# Tretinoin, a Vitamin A derivative, exerts antiviral effects against Chikungunya virus infection through the nuclear retinoid receptor signaling pathways

**DOI:** 10.1101/2025.09.29.679315

**Authors:** Soumyajit Ghosh, Chandan Mahish, Archana Mahapatra, Tathagata Mukherjee, Pratima Kumari, Bijita Bhowmick, Saikat De, Udvas Ghorai, Rajshree Rajmohan Jena, Seema Pradhan, Subhasis Chattopadhyay, Soma Chattopadhyay

## Abstract

Chikungunya virus (CHIKV) has reemerged as a global pathogen causing serious public health threat and socio-economic damage, particularly due to the absence of specific antivirals. Retinoids have been reported to exhibit antiviral potentials against multiple viral infections. In the present study, Tretinoin (TR) or all-trans retinoic acid (ATRA) was evaluated for its anti-CHIKV activity using in *vitro*, *in vivo*, and *ex vivo* models. TR was able to impede CHIKV infection efficiently in Vero and physiologically relevant muscle cells, C2C12, with drastic reduction in the viral RNA, proteins and progeny formation. The IC_50_ of TR against CHIKV was estimated to be 20.05 µM and 19.71 µM in Vero and C2C12 cells, respectively. In addition, the inhibition was effective during entry as well as in the early stages of post CHIKV-infection. Notably, TR showed remarkable virucidal activity. Next, global transcriptome profile demonstrated that the signaling by Retinoic acid pathway was highly upregulated after CHIKV infection and Retinoic acid receptors (RAR), specifically RAR-β are the key mediators for the anti-CHIKV effect of TR. Further, the drug was also capable to modulate CHIKV-induced inflammatory responses by reducing the phosphorylated MAPKs, NF-κB and proinflammatory cytokines. Interestingly, TR protected C57BL/6 mice from CHIKV challenge by diminishing viral burden leading to reduced clinical scores and better survival. Similar observation was also noticed in the *ex vivo* model of CHIKV-infected hPBMC-derived monocyte-macrophage populations. In conclusion, this work demonstrated the repurposing potential of TR against CHIKV infection for the first time, encouraging its clinical validation towards future therapeutics.

**IMPORTANCE:** CHIKV is an arbovirus causing Chikungunya Fever (CHIKF) leading to debilitating arthralgia and myalgia. The unavailability of licensed antivirals or potent vaccines against CHIKV encourages extensive research to find a therapeutic cure. Retinoids, the derivatives of vitamin A, have been repurposed as antiviral agents against a broad range of viruses; however, their efficacy against alphaviruses remains unexplored. Accordingly, Tretinoin (TR), also known as all-trans retinoic acid (ATRA), was validated for anti-CHIKV efficacy in this investigation. Significant perturbation of CHIKV titer through the modulation of Retinoic acid receptors (RAR) was observed upon drug-treatment. Further, TR reduced major inflammatory markers (MAPKs and proinflammatory cytokines) supporting its anti-inflammatory effect. Further, its anti-CHIKV potential was validated in the mouse model and hPBMC-derived monocyte-macrophage cells, indicating the pre-clinical efficacy. Finally, this is the first study to report the anti-CHIKV property of Tretinoin using *in vitro*, *in vivo*, and *ex vivo* approaches suggesting its repurposing potential.

## INTRODUCTION

Chikungunya virus (CHIKV) is a vector-borne Alphavirus belonging to the *Togaviridae* family and transmitted to humans by mosquitoes of the *Aedes* genus. Following its first outbreak in 1952 in the Makonde plateau (Tanzania), CHIKV outbreaks were mostly sporadic across tropical and subtropical regions of Africa, Asia, Oceania, Europe and the Americas (1). Acute CHIKV infection leads to a febrile disease namely Chikungunya Fever (CHIKF) manifesting high fever, debilitating arthralgia (joint pain), myalgia (muscle pain), headache, nausea, asthenia, gastrointestinal distress and maculopapular rashes (2, 3). Even neurological complications have also been reported along with other comorbidities like encephalitis, optic neuropathy, neuroretinitis and Guillain – Barré syndrome (4). Although CHIKF is rarely fatal and symptoms are often self-limiting, it can cause severe morbidity and death in adults with underlying medical conditions (5). The growing emergence of CHIKV infection has raised a major global public health concern, particularly due to the unavailability of licensed drugs, specific antiviral therapies or effective vaccine candidates. The World Health Organization (WHO) has categorized CHIKV infection as a neglected tropical disease.

CHIKV is a spherical (approximately 70 nm diameter), enveloped, single-stranded RNA virus with a positive sense genome of about 11.8 kb, containing two open reading frames (ORFs) (3). The 5′ end of the ORF encodes four non-structural proteins, nsP1-4 (replicase polyprotein) and the 3′ end is translated from a subgenomic promoter encoding the structural proteins [capsid protein (C), glycoproteins: E1, E2, E3, and 6K) which assemble to form progeny virions (6, 7). As no licensed antivirals are available against CHIKV or other alphaviruses till date, there is a critical need to find therapeutic interventions. Non-steroidal anti-inflammatory drugs (NSAIDs), analgesics and antipyretics have been recommended to alleviate the symptoms in CHIKF patients (8). Though several drugs and compounds like chloroquine, ribavirin, arbidol, chlorpromazine, imipramine, mefenamic acid, flavipiravir etc. were repurposed against CHIKV, their therapeutic benefits remain limited (9). A live-attenuated, single dose investigational vaccine candidate, Ixchiq (VLA-1553) has recently been approved by the U.S. FDA in November 2023 (10); however, its clinical efficacy still requires validation through large-scale community-based trials. These necessitate the research to develop potent antiviral candidates having *in vitro* as well as *in vivo* efficacy against CHIKV.

Finding a new drug is prohibitive in terms of cost and time. On the contrary, Drug repurposing is a promising and cost-effective alternative for quicker clinical application (11). Over the past two decades, many studies have documented that vitamin A supplementation leads to significant reduction in mortality in young children in random community-based trials (12, 13). It has also been observed that populations having vitamin A deficiency are highly susceptible to infectious diseases (14). The natural and synthetic derivatives as well as metabolites of vitamin A (Retinol) are collectively referred to as Retinoids, governing a broad range of biological processes including regulation of embryonic development, maintenance of the integrity of epithelial cell surfaces, vision and immunity (15, 16). Retinoic acid (RA) is the biologically active form of Retinol, having three isomers: all-trans retinoic acid (ATRA), 9-cis retinoic acid (9cis-RA), and 13-cis retinoic acid (13cis-RA). Several studies have deciphered the antiviral properties of retinoids against a broad spectrum of virus families, viz. *Retroviridae* (17), *Herpesviridae* (18), *Paramyxoviridae* (19, 20), *Caliciviridae* (21), *Hepadnaviridae* (22, 23), *Flaviviridae* (24–26) and *Coronaviridae* (27, 28). Since there are no such evidences of antiviral effects against alphaviruses by retinoids, the natural and synthetic retinoids can be tangible solutions for the management of CHIKV infection. Among the RAs, Tretinoin (TR), also known as all-trans-retinoic acid (ATRA) is effective for treating skin diseases and preventing acute promyelocytic leukaemia (APL) (29).

The current study aims to evaluate the repurposing potential of ATRA or TR against CHIKV infection for the first time. To elucidate the anti-CHIKV efficacy as well as the underlying mechanism of action of the drug, the investigation was carried out across multiple biological systems: *in vitro* in various cell lines, *in vivo* in C57BL/6 mice, and *ex vivo* using human peripheral blood mononuclear cell (hPBMC)-derived monocyte-macrophage cells.

## RESULTS

### TR abrogates CHIKV infection efficiently

The MTT assay was executed to assess the cytotoxicity of TR in Vero cells. The cell viability was >95% at 100 μM concentration of the drug and 50% cytotoxicity (CC_50_) was observed to be 167.3 μM as shown in Fig. 1A. Thus, a dose of 100 μM or less was used to evaluate its anti-CHIKV efficacy in Vero cells. The cells were infected with CHIKV with a multiplicity of infection (MOI) of 0.1 and treated with different concentrations of TR post-infection. Supernatants and cells were harvested at 18 hours post-infection (hpi) and subjected to plaque assay and immunofluorescence imaging, respectively. The antiviral effect was also very prominent as viral titer was reduced drastically in CHIKV-infected cells upon drug treatment in a dose-dependent manner (Fig. 1B). The IC_50_ of TR was found to be 20.05 μM against CHIKV in Vero cells and the selectivity index (SI) was estimated to be 8.34 (Fig. 1C). Comparing the drug-treated cells to the infection control, a considerable decrease in Cytopathic effect (CPE) was observed (Fig. 1D). Further, the immunofluorescence analysis revealed a gradual decrease in the number of viral E2 positive cells compared to the infection control (Fig. 1E and F). Hence, these findings indicate that TR can inhibit CHIKV infection efficiently.

**Figure 1:**
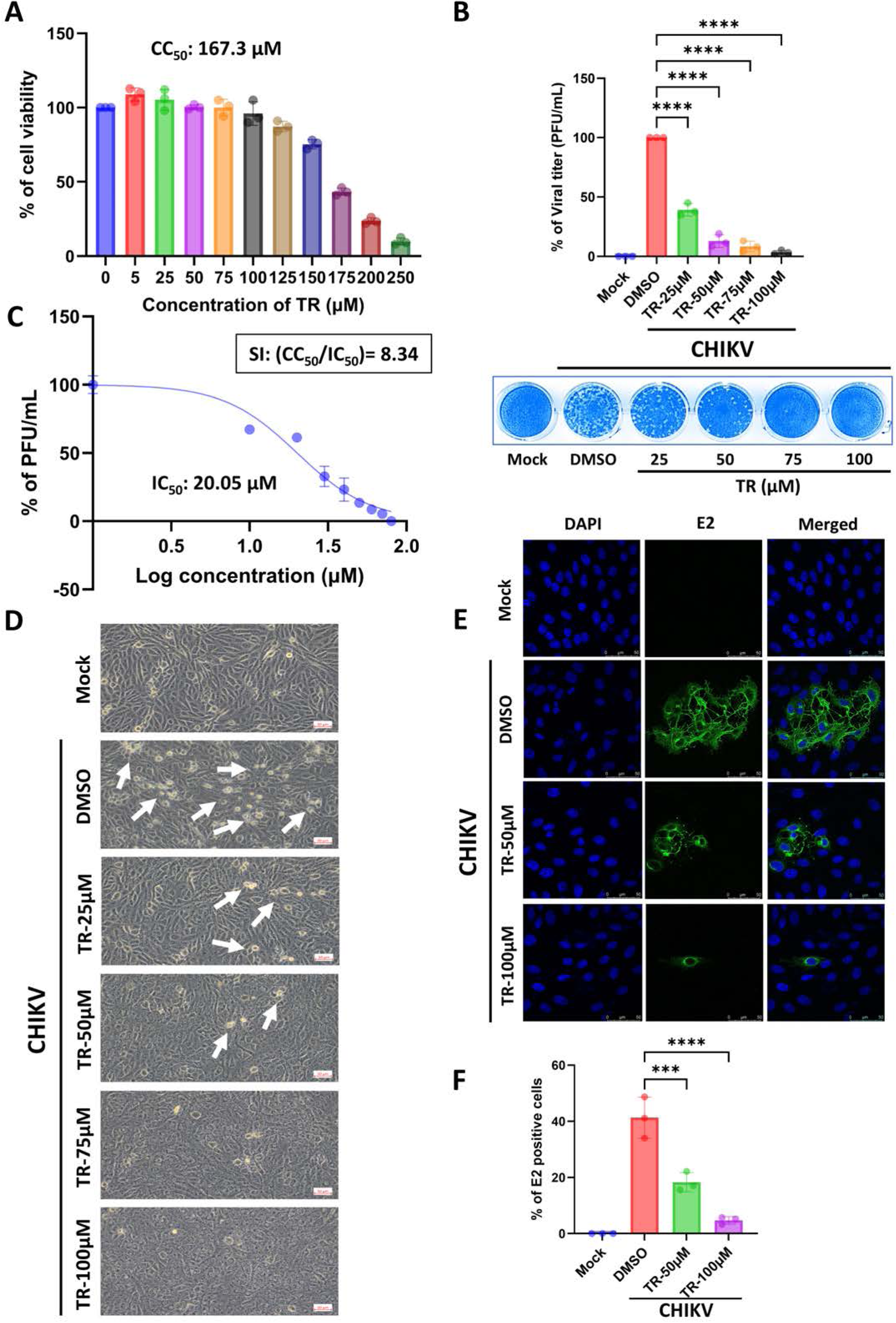
TR inhibits CHIKV infection efficiently. Vero cells were treated with different concentrations of TR, followed by MTT assay to calculate the 50% cytotoxicity (CC_50_). (A) Bar diagram showing the percentage (%) of cell viability in presence of different concentrations of TR. (B) Vero cells were infected with CHIKV and TR was added at different concentrations (25, 50, 75 and 100 μM) post-infection. The supernatants were collected at 18 hpi and viral titer was determined by plaque assay. The bar diagram represents the % of viral titer. The image in the lower panel represents the respective plaque assay plate. (C) The IC_50_ of TR against CHIKV-infected Vero cells was estimated. The X-axis illustrates the logarithmic value of the different concentrations of TR and the Y-axis portrays the % of PFU/mL. (D) Images depicting the CPE of mock, infected and TR-treated cells, observed under a bright field microscope with 20X magnification at 18 hpi. The white arrows indicate the CPE after CHIKV infection. (E) Vero cells plated onto the cover slips were infected with CHIKV and treated subsequently with different concentrations of TR. At 18 hpi, the cells were fixed and probed with CHIKV-E2 antibody followed by staining with anti-mouse Alexa Fluor 488 (green) secondary antibody for confocal imaging. Nuclei were counterstained with DAPI (blue). Scale bar: 50 µM. (F) Bar diagram indicating the percent of E2 positive cells from the confocal images. Data from three independent experiments are represented as mean ± SD; p < 0.05 was considered statistically significant.

### TR lowers viral RNA and protein levels

To validate the inhibitory effect of TR on CHIKV RNA and proteins, qRT-PCR, Western blotting and flow-cytometry based staining had been performed. CHIKV-nsP2 and E1 genes were amplified from total RNA by qRT-PCR from virus-infected and treated cells using gene-specific primers. It was found that the levels of nsP2 and E1 gene expressions had been decreased by 10-fold and 20-fold, respectively, in the presence of 100 μM of TR (Fig. 2A and B). Interestingly, the Western blots demonstrated 80-90% reduction of the non-structural (nsP2) and structural (E2) proteins in the presence of TR (100 μM) (Fig. 2C to E). Next, Flow cytometric dot plot analysis revealed a substantial reduction of CHIKV-E2 positive cells with increasing concentrations of the drug (Fig. 2F). Collectively, the results indicate that TR significantly impedes CHIKV infection by reducing viral RNAs and proteins.

**Figure 2:**
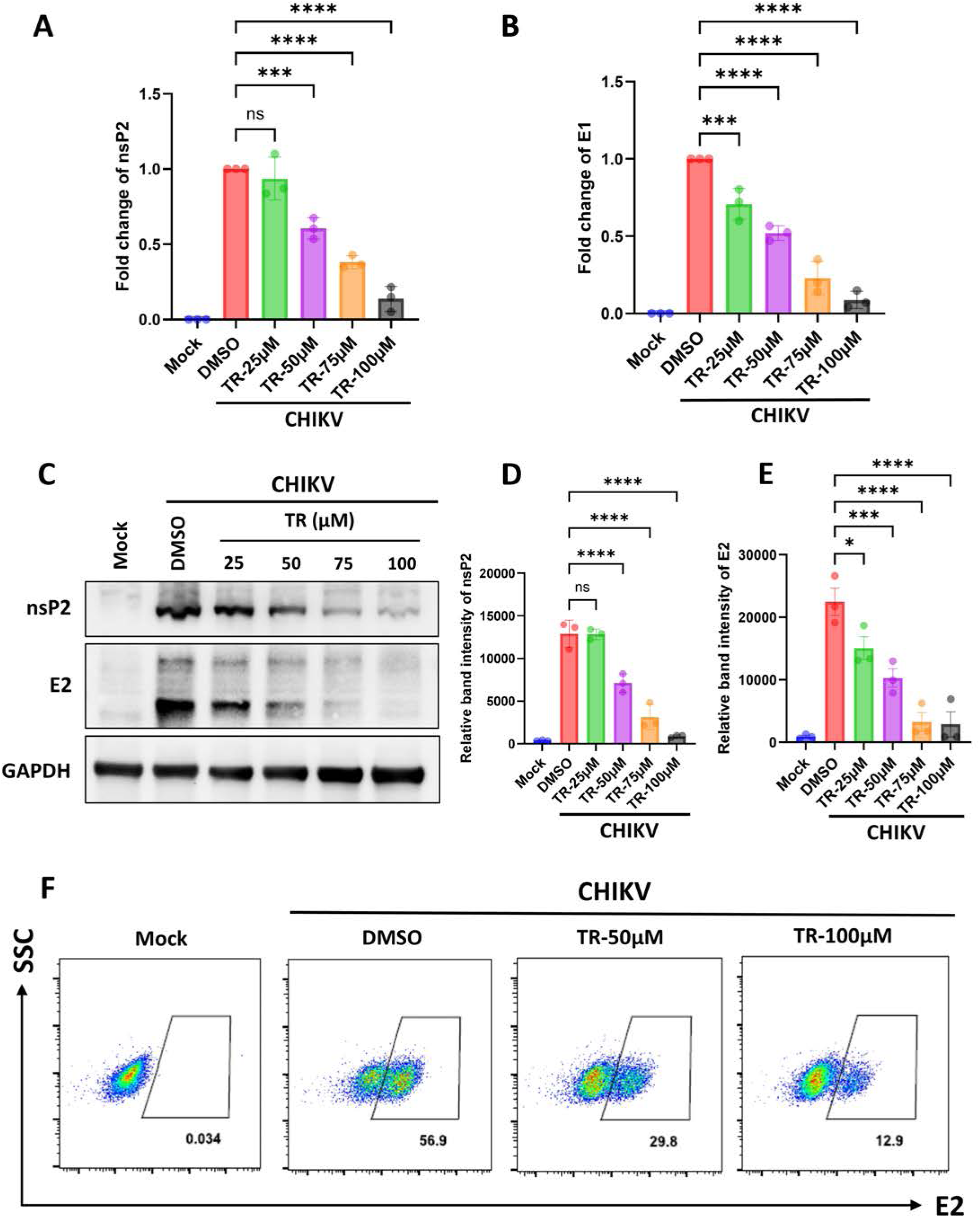
TR reduces viral RNA and protein levels. Vero cells were infected with CHIKV and treated with different concentrations of TR. At 18 hpi, cells were harvested with Trizol reagent and total RNA was extracted from the mock, CHIKV-infected and treated cells and qRT-PCR was performed. The CHIKV-nsP2 and E1 genes were amplified by qRT-PCR. (A, B) Bar diagrams displaying the fold changes of viral nsP2 and E1 genes. (C) Cell lysates were processed for Western blotting using CHIKV-nsP2 and E2 antibodies. GAPDH served as a loading control. (D, E) Bar diagrams showing the relative band intensities of nsP2 and E2 of the Western blot. (F) Flow cytometric dot plot analysis depicting percent cells positive for CHIKV-E2 in presence or absence of TR. Data presented as mean ± SD (*n* = 3; p < 0.05 was considered statistically significant).

### TR attenuates CHIKV infection in the physiologically relevant muscle cells

To evaluate the antiviral efficacy of TR, a physiologically relevant mouse myoblast cell line, i.e., C2C12 was selected. The CC_50_ of TR was found to be 189.6 μM through MTT assay (Fig. 3A). The cells were then infected with CHIKV at 0.05 MOI and treated with different non-toxic doses of TR. Cells and culture supernatants were harvested at 16 hpi and subjected to plaque assay, immunofluorescence imaging, qRT-PCR, Western blotting, and and flow-cytometry based staining. TR was able to impede viral progeny release significantly (60% and >80% reduction by 50 and 100 µM TR, respectively) (Fig. 3B). The IC_50_ of TR in C2C12 cells was estimated to be 19.71 µM against CHIKV and the SI was calculated to be 9.61 (Fig. 3C). Further, a remarkable decrease in the CPE was observed upon addition of TR, compared to the infection control (Fig. 3D). Interestingly, the confocal microscopy exhibited a drastic reduction of CHIKV E2 antigen in presence of TR (Fig. 3E and F). Next, the qRT-PCR data demonstrated 10-fold and 6-fold decrease in nsP2 and E1 gene expressions, respectively upon drug treatment (100 µM) (Fig. 3G and H). Moreover, the Western blots exhibited a dose-dependent depletion of nsP2 and E2 proteins with increasing concentrations of the drug. TR (100 µM) reduced the nsP2 and E2 protein levels by >90% and 85%, respectively (Fig. 3I to K). Additionally, a significant reduction in CHIKV-E2 positive cells was observed by flow-cytometry in presence of the drug (Fig. 3L). Altogether, these findings suggest an effective inhibitory capacity of TR in abrogating CHIKV infection in C2C12 cells.

**Figure 3:**
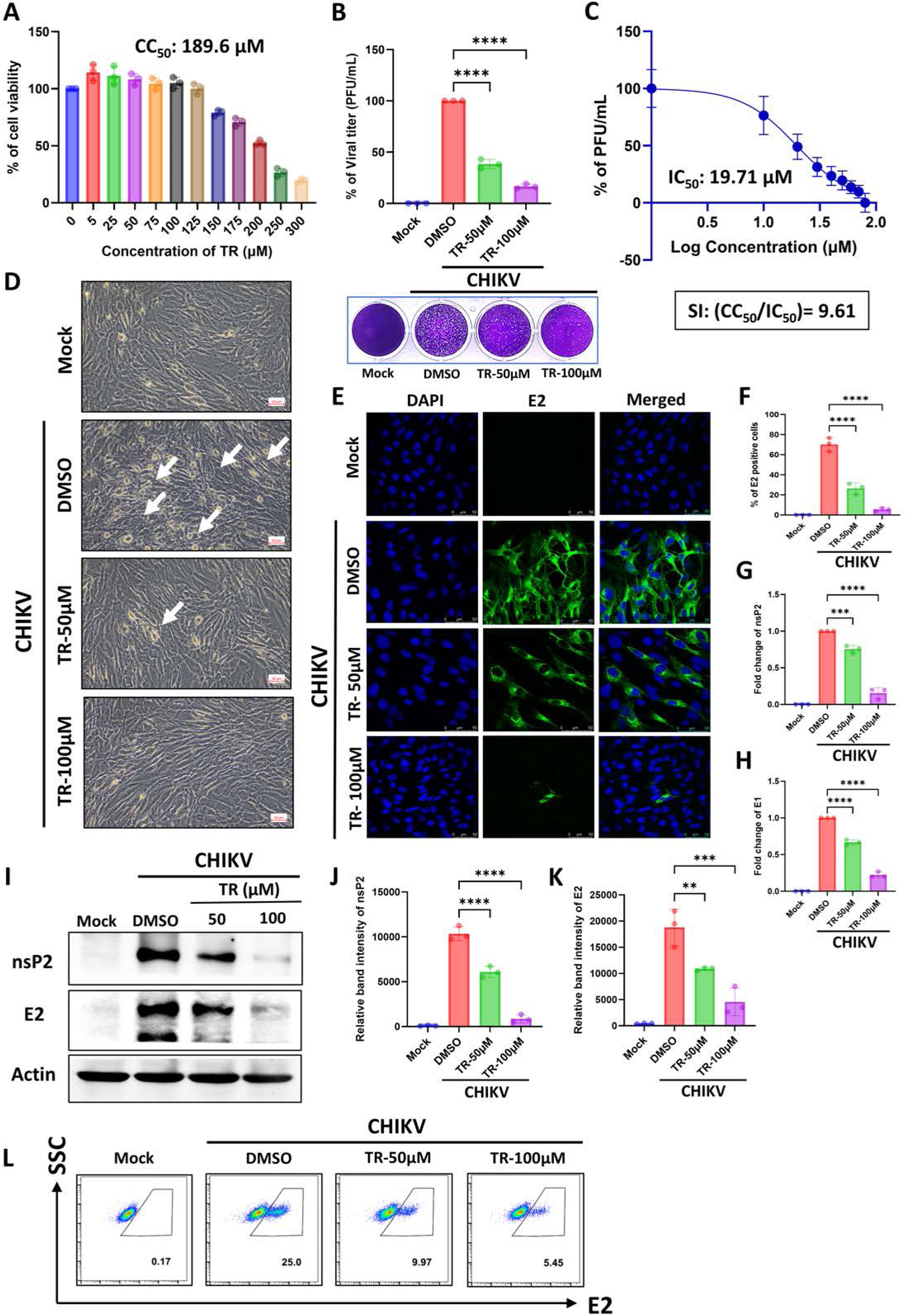
TR abrogates CHIKV infection in the physiologically relevant muscle cells. C2C12 cells were treated with different concentrations of TR and CC_50_ value was estimated through MTT assay. (A) Bar diagram showing the cell viability. (B) Muscle cells were infected with CHIKV and subsequently treated with different non-toxic doses of the drug. The supernatants were collected at 16 hpi for plaque assay. The bar diagram depicts the % of viral titer of supernatants. The lower panel represents the respective plaque assay plate image. (C) The IC_50_ of TR against CHIKV in C2C12 cell was calculated. The X-axis represents the logarithmic value of different concentrations of TR and the Y-axis illustrates the % of PFU/mL. (D) At 18 hpi, CPE was observed under a bright field microscope and images of the mock, infected and drug-treated cells were taken with 20X magnification. Arrows indicating the CPE after CHIKV infection. (E) Confocal images of Mock, infected or drug-treated muscle cells which were probed with CHIKV-E2 antibody. Nuclei were counterstained with DAPI. Scale bar: 50 μM. (F) Bar diagram exhibiting the percent positive E2 cell counts from the confocal images. (G, H) Total RNA was isolated from the mock, CHIKV-infected and drug-treated cells. The CHIKV-nsP2 and E1 genes were amplified by qRT-PCR. Bar diagrams indicating the fold changes of nsP2 and E1 genes. (I) Western blot images showing nsP2 and E2 levels of mock, infected and TR-treated cells. Actin was used as a loading control. (J, K) Bar diagrams displaying the relative band intensities of viral nsP2 and E2 proteins. (L) Flow cytometry-based dot plot analysis showing % positive cells for CHIKV-E2 after infection and drug treatment. All the experiments were carried out in triplicate and data presented as mean ± SD; p < 0.05 was considered statistically significant.

### TR shows anti-CHIKV efficacy during entry and post-entry phases of infection and affects the early stages of CHIKV life cycle post-infection

In order to understand the phase of the viral life cycle which is abrogated by the drug, TR (50 μM) was administered at different phases of CHIKV infection (pre, during and post treatment) in both Vero and C2C12 cell lines. The findings revealed that pre-treatment with TR had no inhibition on viral particle formation whereas during-treatment led to 60% reduction in viral progeny formation in the Vero and 40% in the C2C12 cells (Fig. 4A and B). Nonetheless, the best inhibitory potential was found upon the post-treatment condition where the viral titer was abrogated by 85% in Vero and 75% in C2C12 cells (Fig. 4A and B). Next, to evaluate the extracellular effect of the drug on CHIKV particles, virucidal assay was performed. Interestingly, the virucidal assay resulted in a drastic reduction (∼80%) of viral titer in the TR-treated sample as compared to untreated control (Fig. 4C). Further, to elucidate the stage of CHIKV life cycle post-infection, where the drug is effective, a “time-of-addition” experiment was performed. The release of infectious virus particles was abrogated with the addition of TR (50 µM) at 0 to 8 hpi by 15% to 70% in Vero and 18% to 60% in case of C2C12 cells, compared to the infection control (Fig. 4D and E). However, the inhibition was not effective (from 10% to 0%) when drug was added in the late stages between 8 to 16 hpi (for Vero) or 10 to 14 hpi (for C2C12). Taken together, it can be suggested that TR is effective in the during and post-treatment conditions and might interfere in the early stages of CHIKV life cycle post-infection.

**Figure 4:**
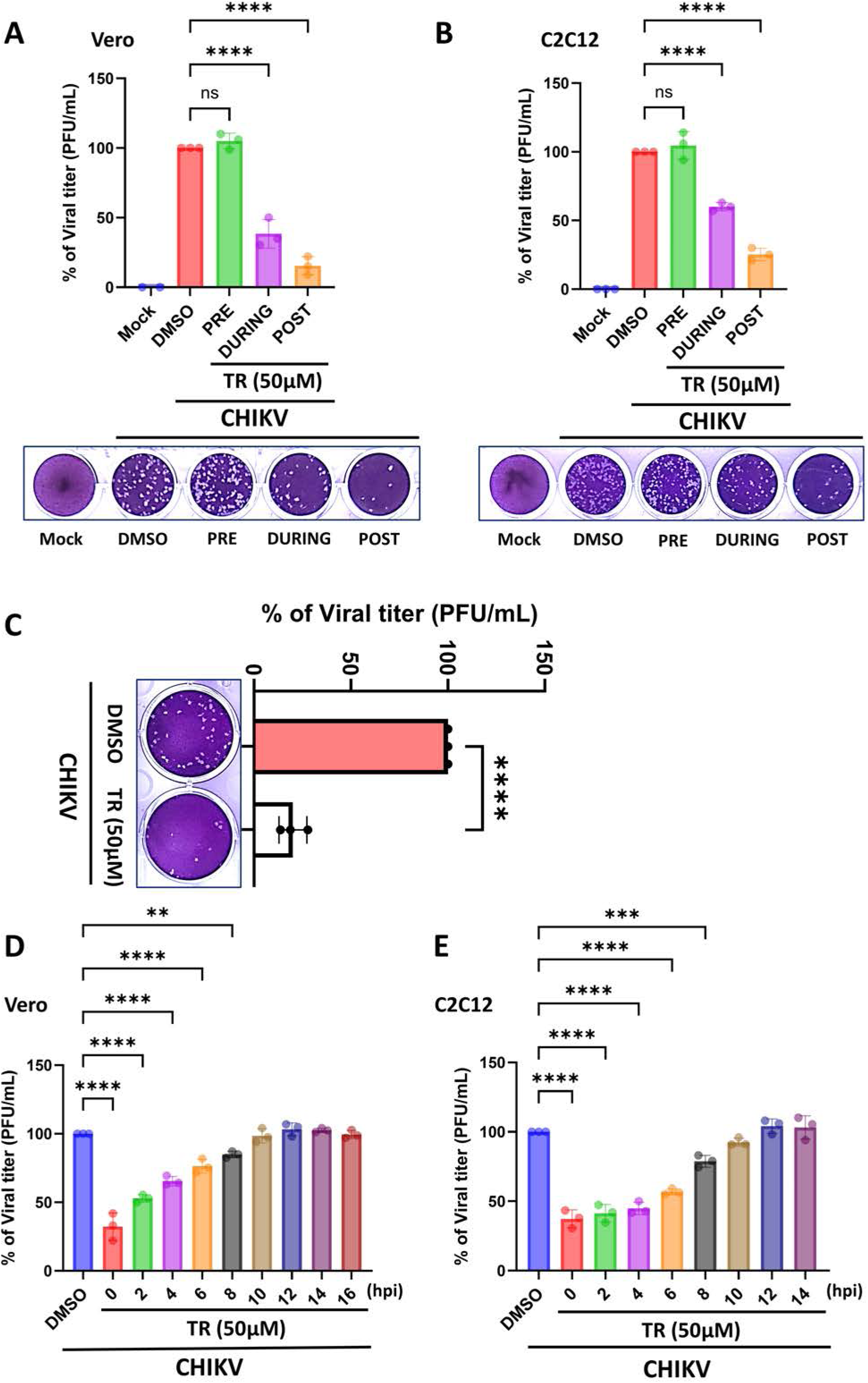
TR is effective during entry and post-entry early stages of CHIKV infection. Vero and C2C12 cells were treated with TR (50 µM) separately in three different conditions (pre, during and post infection). Supernatants were collected at 18 hpi for Vero and 16 hpi for C2C12 cells and subjected to plaque assay. (A, B) The bar diagrams indicate the % of viral titer of the supernatants collected after different treatment conditions. The images in the lower panel demonstrate the corresponding plaque assay plates. (C) Bar diagram depicting the % of CHIKV titer after performing the virucidal assay. Left panel represents the image of respective plaque assay plate. (D, E) Both the Vero and C2C12 cells were infected with CHIKV and 50 µM drug was added to the cells at every 2 h intervals up to 14 or 16 hpi (16 hpi for Vero and 14 hpi in case of C2C12 cells). Bar diagrams portray the % of viral titer, obtained by plaque assay from all the supernatant samples collected at 16 or 18 hpi. Data of three independent experiments are shown as mean ± SD; p < 0.05 was considered statistically significant.

### TR alters expression of the genes associated to nuclear retinoic acid receptors

In order to decipher the mechanism of action of TR against CHIKV infection, the impact of the drug on global gene expression profiles in both uninfected (mock) and infected C2C12 cells was investigated by RNA Sequencing (RNA-seq) (Fig. 5A). Analysis of the differentially expressed genes (DEGs) revealed distinct transcriptional signatures of the mock, infected, mock-treated and/or infected-treated cells (Venn diagram, Fig. S1A) indicating the extreme coordinates on the Principal Component Analysis (PCA) plot (Fig. 5B). Further, Reactome and Gene Set Enrichment Analysis (GSEA) identified a series of cellular pathways to be significantly modulated after CHIKV infection and drug treatment. Notably, a similar pattern of transcriptional changes was observed upon addition of TR to both the mock and infected cells, as illustrated by the expression cluster heatmap comparing Mock and Mock + TR conditions (Fig. S1B). Next, Reactome analysis comparing the uninfected and infected conditions (Mock vs. CHIKV) identified the ‘Signaling by Retinoic acid’ pathway to be highly upregulated upon CHIKV infection (Fig. 5C). Since TR is a class of Retinoic acid, its effect on the genes associated with the Retinoic acid signaling pathway was analyzed further. Among them, Retinoic acid receptor gamma (RAR-γ) and some metabolic genes, such as cytochrome P450 family 26 subfamily C member 1 (CYP26C1), carnitine palmitoyltransferase 1B (CPT1B), fatty acid binding protein 5 (Fabp5) and dehydrogenase/reductase 3 (DHRS3) were the most significant DEGs (Fig. 5D). In addition, GSEA between infected and drug-treated condition (CHIKV vs CHIKV+TR) revealed upregulation of several biological pathways, viz. Transport of small molecules, Drug ADME, Biological oxidations, Signaling by Nuclear Receptors, Phase II − Conjugation of compounds, Neutrophil degranulation, Cellular response to chemical stress and Innate Immune System (Fig. 5E). Interestingly, the retinoic acid receptors (RARs), belong to the Nuclear Receptor family, are activated by retinoic acid and form heterodimers with retinoid X receptors (RXRs) (30). Thus, the expression profiles of the RAR-related genes were assessed further on the basis of Z-score analysis across different sample groups. RAR-β and RAR-γ were found to be significantly modulated upon drug treatment as compared to infection (Fig. 5F). Moreover, protein–protein interaction analysis using the STRING database showed strong interactions between RAR-β and RAR-γ (Fig. 5G). Collectively, these findings suggest that TR might exert its anti-CHIKV effect via the modulation of nuclear retinoid receptor signaling, partly through RAR-β and RAR-γ.

**Figure 5:**
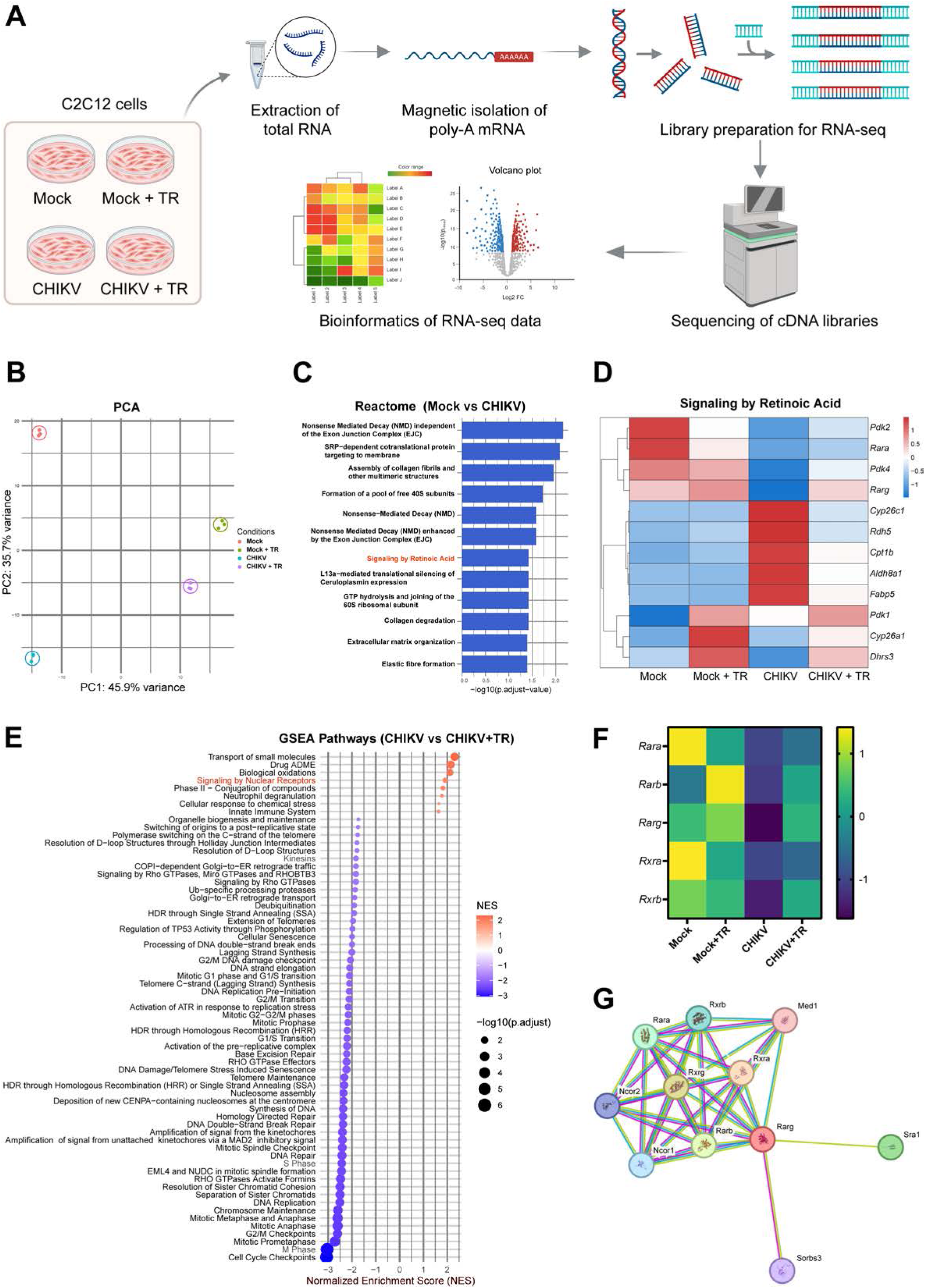
TR alters expression of the genes associated to nuclear retinoic acid receptors. C2C12 cells were infected with CHIKV at 0.05 MOI and TR (100 μM) was added post-infection. Mock and mock-treated cells were taken as control. Cells were harvested at 16 hpi using Trizol reagent for extraction of total RNA, followed by mRNA enrichment, library preparation, sequencing and comparative transcriptome analysis. (A) Schematic workflow of library preparation for the RNA-Seq. (B) PCA plot for the global gene expression profile of the mock, infected, mock-treated and/or infected-treated cells. (C) Reactome Pathway enrichment indicating significantly enriched pathways in Mock vs CHIKV infected condition. (D) Heatmap showing the DEGs enriched for the Signaling by Retinoic acid pathway from the RNA-seq data. (E) Upregulated and downregulated pathways in CHIKV infected vs infected-treated condition, obtained by GSEA. (F) Heatmap demonstrating expression profile of RAR-related genes on the basis of mean Z scores from RNA-seq results. (G) Interaction network of RARγ, obtained from the STRING database. Data shown in figure is combined from three independent experiments; p < 0.05 was considered statistically significant.

### The anti-CHIKV effect of TR is mediated partially through RAR-β

The biological activity of the retinoic acid is exhibited through different RARs. Accordingly, the gene expression levels of RAR-α, β and γ were assessed by qRT-PCR in C2C12 cells under different conditions (Mock, Mock + TR, CHIKV and CHIKV + TR). The qRT-PCR data exhibited non-significant change of RAR-α expression across all the four conditions (Fig. 6A). In contrast, RAR-β gene expression was decreased by 1.5-fold during infection, but upregulated by 1.6-fold and 2-fold in the uninfected and infected cells, respectively upon drug-treatment, compared to the mock control (Fig. 6B). In addition, RAR-γ expression was not altered upon addition of TR to the mock cells but exhibited a 3-fold increase in the infected as well as treated cells, compared to both mock and infected conditions (Fig. 6C). Further, to find out the involvement of specific RAR through which TR inhibits CHIKV, RAR-α specific antagonist RO41-5253 (RO), RAR-β specific antagonist LE135 (LE) and RAR-γ specific antagonist LY-2955303 (LY) were used in this study. MTT assay revealed that RO, LE and LY were non-toxic to C2C12 cells at 50 µM concentration (Fig. S2A, C and E). Treatment with these antagonists did not affect respective RAR-α, β or γ at their RNA expression levels (Fig. S2B, D and F). Next, C2C12 cells were pre-treated separately with the inhibitors 8 h prior to infection, followed by CHIKV infection and TR treatment. The supernatants were collected at 16 hpi and subjected to plaque assay and qRT-PCR. TR treatment alone (20 µM) led to ∼50% reduction and drug treatment in combination with pre-incubation of RO or LY (50 µM) resulted in 70–75% decrease in CHIKV titer (Fig. 6D). However, pre-treatment with LE (50 µM) partially antagonized the antiviral effect of TR as viral titer was reduced only by ∼10% in this case (Fig. 6D). These observations were further validated by qRT-PCR from the viral RNA, which revealed significant decrease in CHIKV RNA copy number upon drug treatment separately (40%) as well as in combination of pre-treatment conditions with RO or LY (65-70%) (Fig. 6E). But TR-mediated reduction in viral RNA was not significant when cells were pre-treated with LE (Fig. 6E). Taken together, these results indicate the potential role of RAR-β in mediating the anti-CHIKV effect of TR.

**Figure 6:**
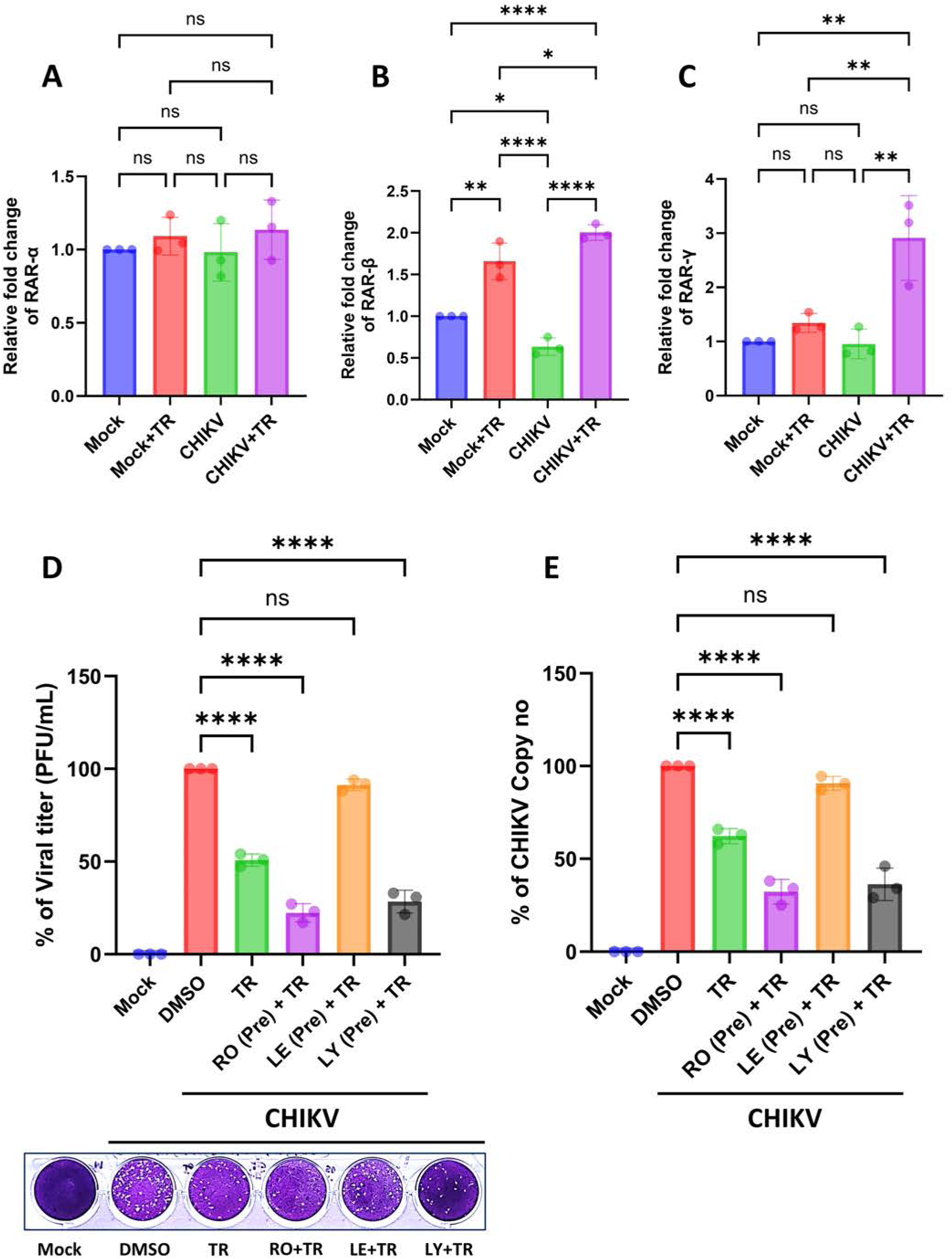
The anti-CHIKV effect of TR is mediated partially through RAR-β. C2C12 cells were infected with CHIKV and treated subsequently with TR (100 μM). Cells were harvested at 16 hpi using Trizol reagent. Total RNA was extracted from the mock, infected, mock-treated and infected-treated cells and the RAR-α, β and γ genes were amplified by qRT-PCR. (A to C) Bar diagrams displaying the fold changes of RAR-α, RAR-β and RAR-γ under four different conditions, viz. Mock, Mock + TR, CHIKV and CHIKV + TR. (D) C2C12 cells were pre-incubated with RO, LE and LY (50 µM) for 8 h. Next, the cells were washed with 1X PBS and infected with CHIKV (0.05 MOI) for 90 min and TR (20 µM) was added post-infection. The supernatants were collected at 16 hpi and subjected to plaque assay. Bar diagram represents the % of viral titer from all the supernatants. The lower panel depicts the image of respective plaque assay plate. (E) Viral RNA was isolated from the culture supernatants, followed by cDNA synthesis and then CHIKV-E1 gene was amplified by qRT-PCR. CHIKV RNA copy number was calculated from a standard curve of C_t_ value. The bar diagram portrays the % of CHIKV RNA copy number/mL in the supernatants. The data are represented as mean ± SD (*n* = 3; p < 0.05 was considered statistically significant).

### TR reduces the CHIKV-induced activation of MAPKs, NF-kB and proinflammatory cytokines in RAW 264.7 cells

Studies have shown that Retinoic acid Inducible Gene I (RIG-I), a major retinoid-responsive gene mediates the antiviral action of retinoids against Measles Virus (MeV) and Mumps virus (MuV) (20, 31). However, the inhibition of CHIKV by TR was found to be RIG-I independent by using Huh7.5 cells with nonfunctional RIG-I (32). TR was able reduce CHIKV titer and protein levels significantly in both Huh7 and Huh7.5 cells (Fig. S3B and C). It was further validated by si-RNA based knock down of RIG-I in the Huh7 cells (Fig. S3D to F). Interestingly, Mitogen-activated protein kinase (MAPK) signaling pathway is activated during CHIKV infection (33, 34). To investigate whether TR can modulate CHIKV-induced inflammation, RAW 264.7 cells were infected with CHIKV-IS at an MOI of 5, followed by drug treatment. Next, the cell lysates were harvested at 8 hpi and the activation (phosphorylation status) of p38, extracellular signal-regulated kinase 1/2 (ERK1/2), Stress-activated protein kinase/Jun amino-terminal (SAPK/JNK), nuclear factor κ-light-chain-enhancer of activated B cells (NF-κB) and interferon regulatory factor 3 (IRF3) were estimated. CHIKV infection led to 2 to 3-fold induction in the in the phosphorylation of each of the major mitogen-activated protein kinases (MAPKs): p38, ERK1/2, and SAPK/JNK. However, drug-treatment (100 µM) resulted in significant reduction in the phosphorylation status of the MAPKs [P38 (10-fold), ERK1/2 (2-fold) and SAPK/JNK (2.1-fold)] (Fig. 7A, D, E and F). Further, induced p-NF-κB (p65) level was lowered by 3-fold in presence of TR (100 µM) (Fig. 7G). Similarly, TR (100 µM) also mitigated the activation of the key transcription factors p-IRF3 by 3.1-fold, compared to the infection control (Fig. 7H). Further, CHIKV infection in RAW 264.7 cells led to significant elevation of different proinflammatory cytokines such as tumor necrosis factor-alpha (TNF-α), interleukin-6 (IL-6) and monocyte chemoattractant protein-1 (MCP-1). Interestingly, the elevated levels of the cytokines were also significantly lowered by the drug [TNF-α (20%), IL-6 (60%) and MCP-1 (30%)] (Fig. 7I to K). Altogether, these findings imply that the anti-CHIKV property of TR is partly mediated through the MAPK inflammatory axis. Moreover, reduction of p-NF-κB, p-IRF3 and proinflammatory cytokine levels support the anti-inflammatory effect of TR.

**Figure 7:**
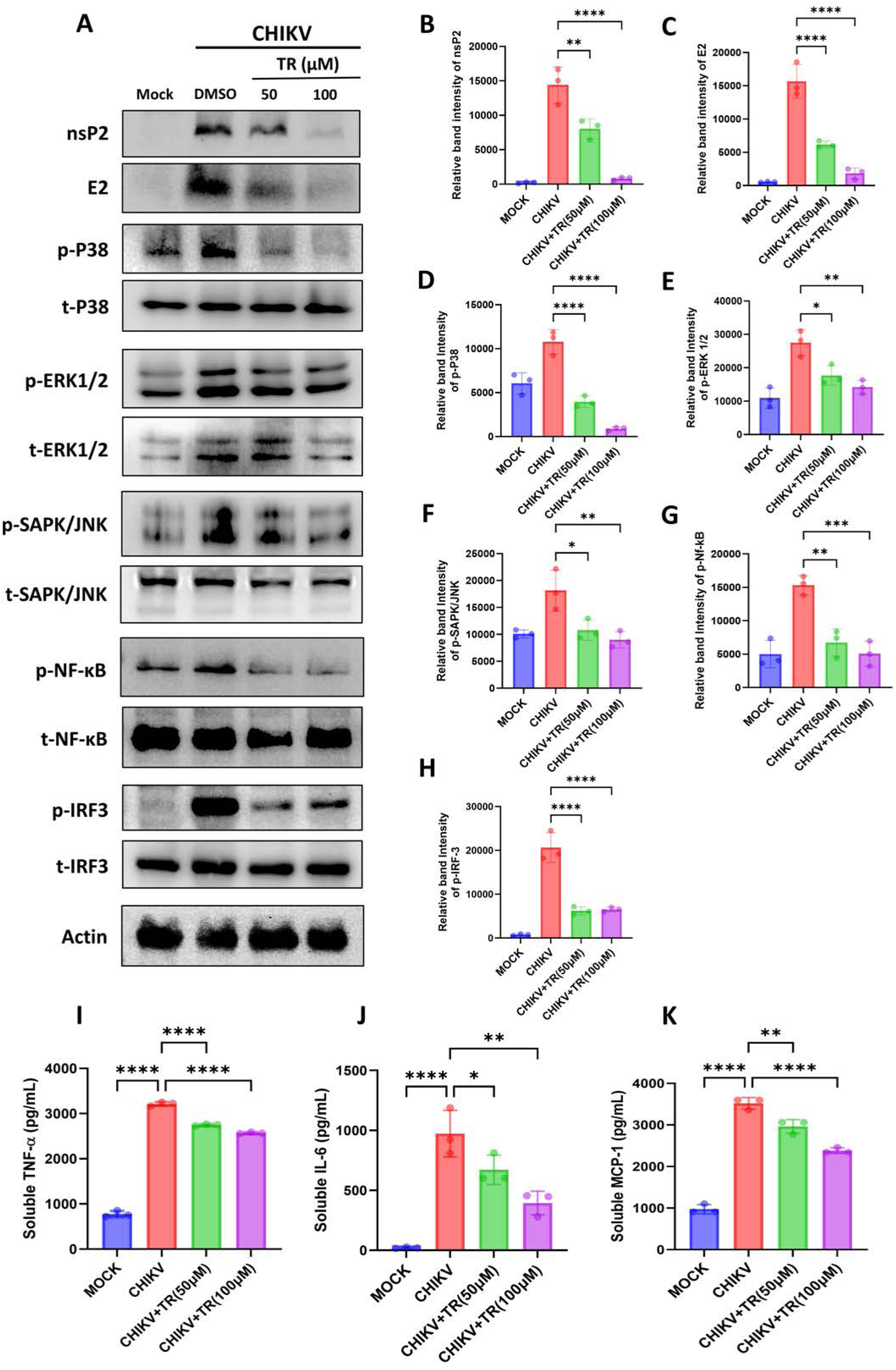
TR lowers (CHIKV-induced) upregulated MAPKs, NF-κB, and proinflammatory cytokines. RAW 264.7 cells were infected with CHIKV, treated with two different doses of TR (50 and 100 μM) post infection and harvested at 8 hpi. (A) The Western blot images exhibiting the nsP2 and E2 protein levels along with phosphorylation status of P38, ERK1/2, SAPK/JNK, NF-κB and IRF3 of mock, infected and drug-treated cell lysates. (B to H) Bar diagrams depicting the relative band intensities of nsP2, E2, p-P38, p-ERK1/2, p-SAPK/JNK, p-NF-κB and p-IRF3. Actin served as a loading control. (I to K) Bar diagrams showing the levels of secreted cytokines (TNF-α, IL-6 and MCP-1) from the supernatants of CHIKV-infected and TR-treated macrophages, quantified using sandwich ELISA. Data of three independent experiments are shown as mean ± SD; p < 0.05 was considered statistically significant.

### TR protects CHIKV-infected mice by diminishing viral burden

In order to validate the anti-CHIKV potential of the drug in the *in vivo* mice model, 10-12 days old C57BL/6 mice were infected subcutaneously with 10^6^ PFU of CHIKV and orally given 5 mg/kg of TR, at every 24 h intervals up to 4 days post infection (dpi) (Fig. 8A). The infected mice group showed CHIKV-induced arthritogenic symptoms, gradual hind limb paralysis and mortality (indicated with red arrows in the figure), while the treated animals showed no such abnormalities leading to reduced clinical score (Fig. 8B). The infected and drug-treated mice were sacrificed on 5 dpi, followed by collection of serum and different tissues for further downstream experiments. Next, plaque assay was performed using equal amount of homogenous and filtered tissues samples (Quadriceps muscle and brain), demonstrating that viral load was reduced by 80% and 70% in the muscle and brain, respectively in the treated mice (Fig. 8C and D). Next, viral RNA was isolated from mice serum, followed by qRT-PCR. Notably, the CHIKV RNA copy number from the mice serum was reduced by ∼50% upon drug treatment (Fig. 8E). In addition, the mice serum samples were subjected to ELISA to estimate the TNF-α levels. Interestingly, TNF-α was found to be significantly less (∼65%) in the treated group, compared to the infected controls (Fig. 8F). In addition, the Western blot analysis from the homogenized tissues revealed significant depletion of nsP2 (90% in the muscle, 70% in the brain) and E2 protein levels (85% and 50% respectively, in the muscle and brain) (Fig. 8G and H). Further, to determine the effect of the drug on the overall disease outcome and survivability, treated or untreated mice infected with CHIKV were monitored for the development of disease symptoms and mortality up to 10 dpi. Infected animals showed severe disease score and all the mice died on 8 dpi. Interestingly, drug treatment minimized the clinical score significantly by alleviating the arthritogenic symptoms (Fig. 8I). Moreover, a dose of 5 mg/kg TR (once per day) provided 70% survival to the CHIKV-infected mice (Fig. 8J). These results suggest the anti-CHIKV efficacy of TR in mice model by impeding viral burden and disease conditions and ensuring better survival against CHIKV infection.

**Figure 8:**
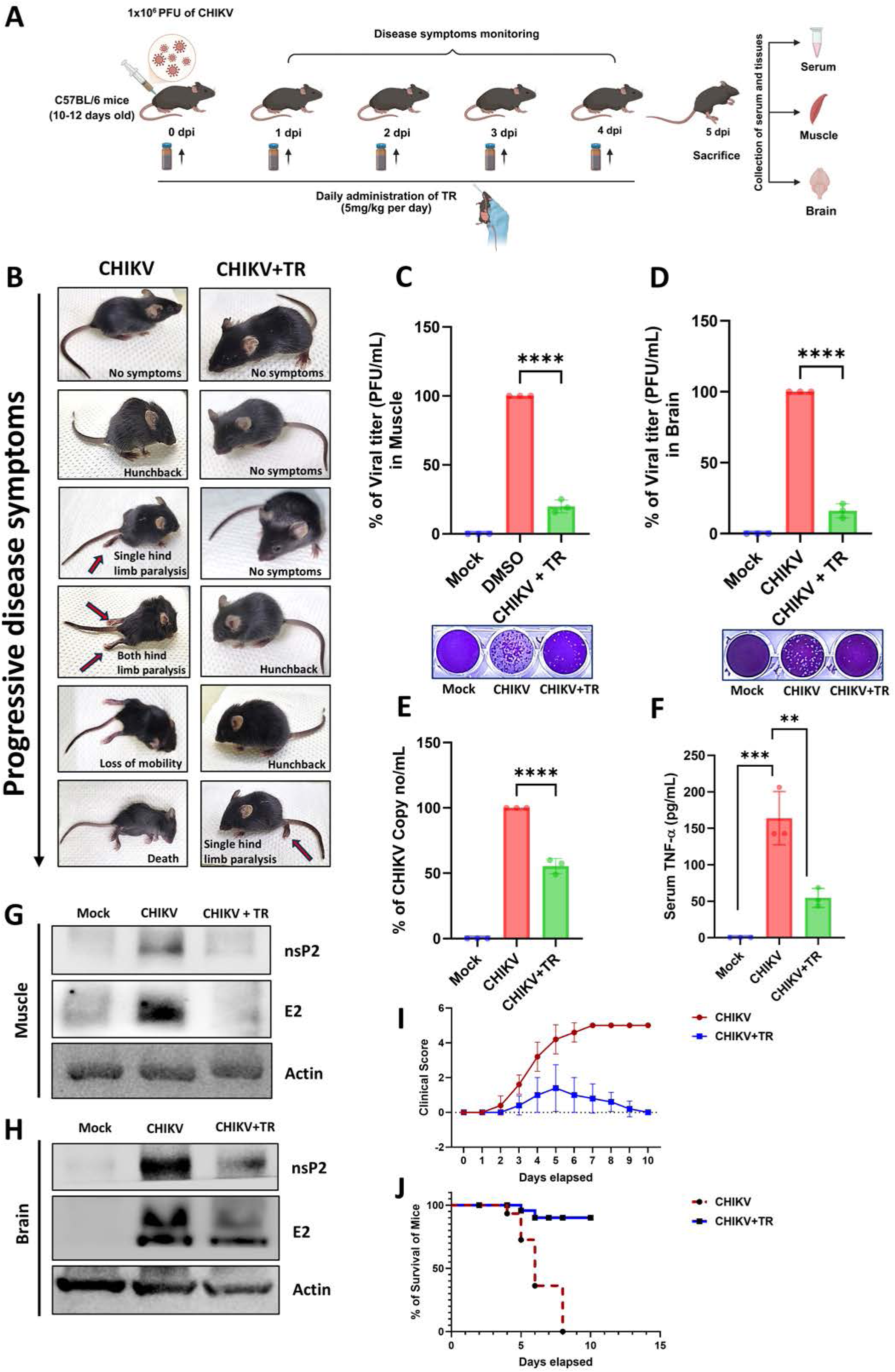
TR protects CHIKV-infected mice by diminishing viral burden. C57BL/6 mice were infected subcutaneously with 10^6^ PFU of CHIKV and treated with 5 mg/kg of TR at every 24 h intervals up to 4 dpi. The animals were sacrificed at 5 dpi; sera, muscle and brain tissues were collected for further downstream experiments. (A) Schematic illustration of the *in vivo* experiment with TR. (B) Images showing CHIKV infected and drug treated mice with symptoms of disease progression (no symptoms: 0; hunchback: 1; single hind limb paralysis: 2; both hind limb paralysis: 3; loss of mobility: 4 and Death: 5). (C, D) Plaque assay was performed to quantitate viral titre using homogenous and filtered tissues samples. Bar diagrams indicating the % of viral titers from infected and drug treated mice muscle and brain, respectively. The lower panel represents the images of respective plaque assay plates. (E) Equal volume of mice serum was taken from mock, infected and drug-treated mice to isolate viral RNA. cDNA was synthesized from equal volume of viral RNA, and the CHIKV-E1 gene was amplified by qRT-PCR. Bar diagram indicating the % of CHIKV RNA copy number/mL in the mice serum. (F) Bar diagram showing the concentration of TNF-α (pg/mL) in mock, infected and treated mice serum. (G, H) Western blots showing the viral nsP2 and E2 proteins in muscle and brain tissue samples. Actin was used as a loading control. (I) Line diagram depicting the clinical scores of the disease symptoms of treated and untreated mice after CHIKV challenge (1 to 10 dpi; *n* = 6/group). (J) Survival curve showing the efficacy of TR against CHIKV-infected C57BL/6 mice (*n* = 6/group). The data are represented as mean ± SD (n=3; p < 0.05 was considered statistically significant).

### TR improves Muscle-histopathology in CHIKV-infected mice

CHIKV infection induces Chronic Skeletal Muscle Atrophy towards disease development in mice (35, 36). Thus, to check whether TR can improve this scenario, histopathological investigations were carried out from the hind limb quadriceps muscles of CHIKV infected and drug treated or untreated mice. The Haematoxylin and Eosin (H&E) staining revealed muscle necrosis as well as acute infiltration of mononuclear lymphocytes in the infected muscle section, which was significantly lowered upon drug treatment (Fig. 9A). Further, immunohistochemistry (IHC) analysis showed significant reduction (∼70%) of CHIKV-E2 antigen in the TR-treated sample, compared to the infected one (Fig. 9B and C). Furthermore, the infiltration status of murine macrophages in the muscles of CHIKV infected and untreated or treated mice were assessed using F4/80 monoclonal antibody. The IHC data demonstrated an intense signal of F4/80 in the infected section which was remarkably decreased by 72% after drug treatment (Fig. 9D and E). Taken together, it can be suggested that TR administration is able to reduce CHIKV induced muscle necrosis and related symptoms *in vivo*.

**Figure 9:**
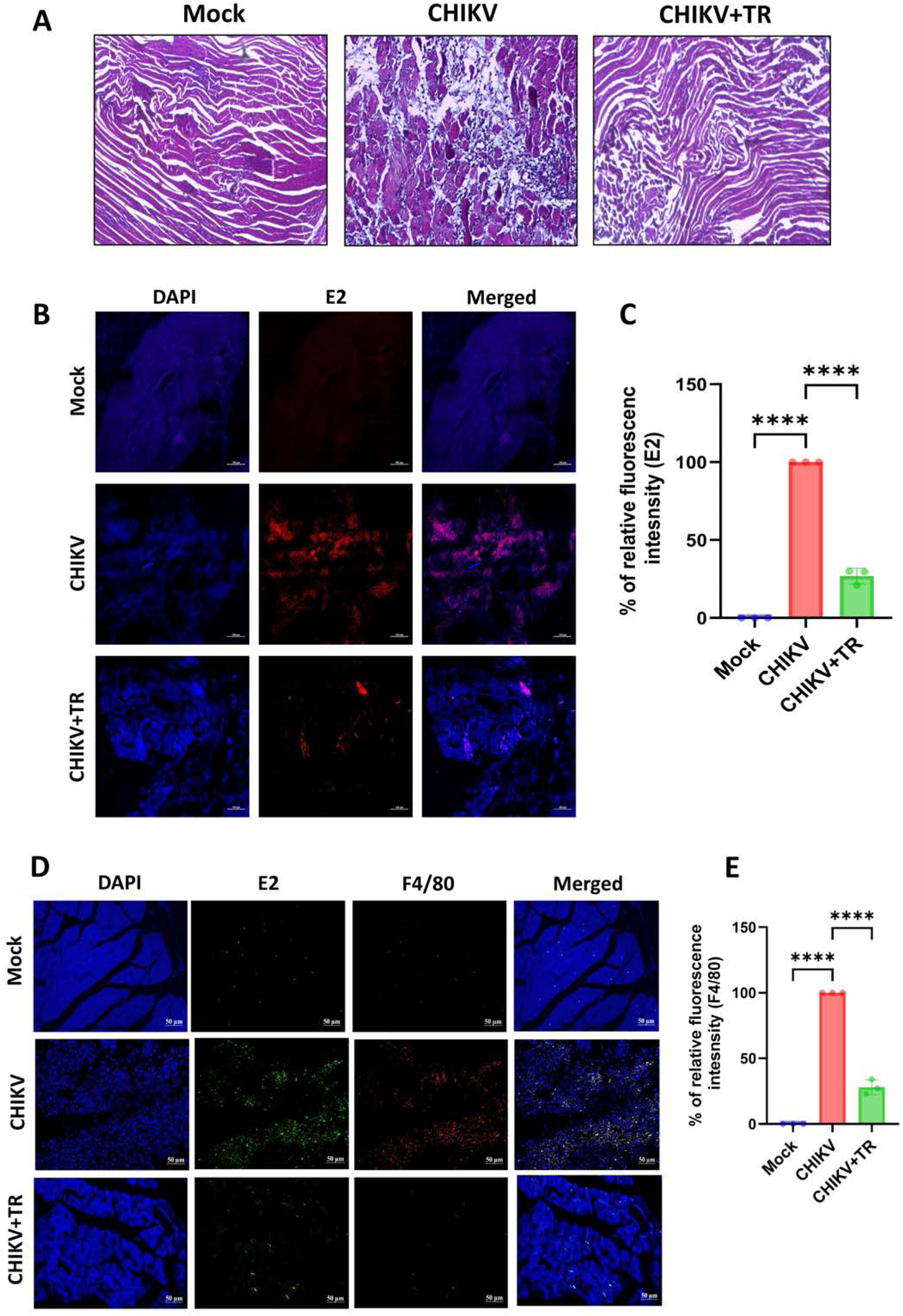
TR improves Muscle-histopathology in CHIKV-infected mice. 10-12 days old C57BL/6 mice were infected subcutaneously with 10^6^ PFU of CHIKV and treated with 5 mg/kg of TR at every 24 h intervals up to 4 dpi. The mice were sacrificed at 5 dpi and hind limb quadriceps muscles were collected for histopathological analysis. (A) Image panels showing H&E-stained muscle sections of uninfected, CHIKV infected and TR-treated animals. (B) Images depicting CHIKV-E2 stained muscle sections, obtained by immunohistochemical analysis (Scale bar: 100 µM). (C) Bar diagram exhibiting the % of relative fluorescence intensity for CHIKV-E2 antigen in different samples. (D) Histological sections of mouse muscles were counterstained with F4/80 monoclonal antibody. CHIKV-E2 antibody was taken as a positive control for infection. The image panels showing the infiltration of murine macrophages in the muscles of mock, infected and treated mice (Scale bar: 50 µM). (E) Bar diagram representing the % of relative fluorescence intensity of F4/80, obtained by the ImageJ software. Data from three independent experiments are shown as mean ± SD; p < 0.05 was considered statistically significant.

### TR perturbs CHIKV infection in the hPBMC-derived monocyte-macrophage populations *ex vivo*

To evaluate the anti-CHIKV efficacy of the drug in higher-order mammalian systems, human peripheral blood mononuclear cells (enriched with CD14^+^/CD11b^+^ cell populations) were used. To characterize the hPBMC-derived adherent cell populations immunologically, B cell (CD19), T cell (CD3), and monocyte-macrophage cell (CD11b and CD14) specific markers were used for flow cytometry. It was observed that the adherent population was predominantly enriched with CD14 + CD11b + monocyte-macrophage cells (Fig. 10A). Next, MTT assay was performed with different concentrations of TR. The result showed the drug is nontoxic even at 250 μM concentration to the CD14 + /CD11b + cells (Fig. 10B). Further, the hPBMC-derived monocyte-macrophage cells collected from three healthy individuals, were infected with CHIKV-IS (MOI: 5) and treated with TR (100 μM). The cells and culture supernatants were harvested at 12 hpi and subjected to flow-cytometry based staining and plaque assay. Interestingly, the CHIKV infected CD14^+^/CD11b^+^ populations showed 31.26% E2-positive cells, whereas treatment with TR resulted in decrease of the E2-positve cells to 22.3% (Fig. 10C and D). In addition, the plaque assay revealed that viral particle formation was reduced by ∼50% upon drug treatment (Fig. 10E). Collectively, these observations indicate that TR can abrogate CHIKV infection significantly in the hPBMC-derived monocyte-macrophage populations *ex vivo*.

**Figure 10:**
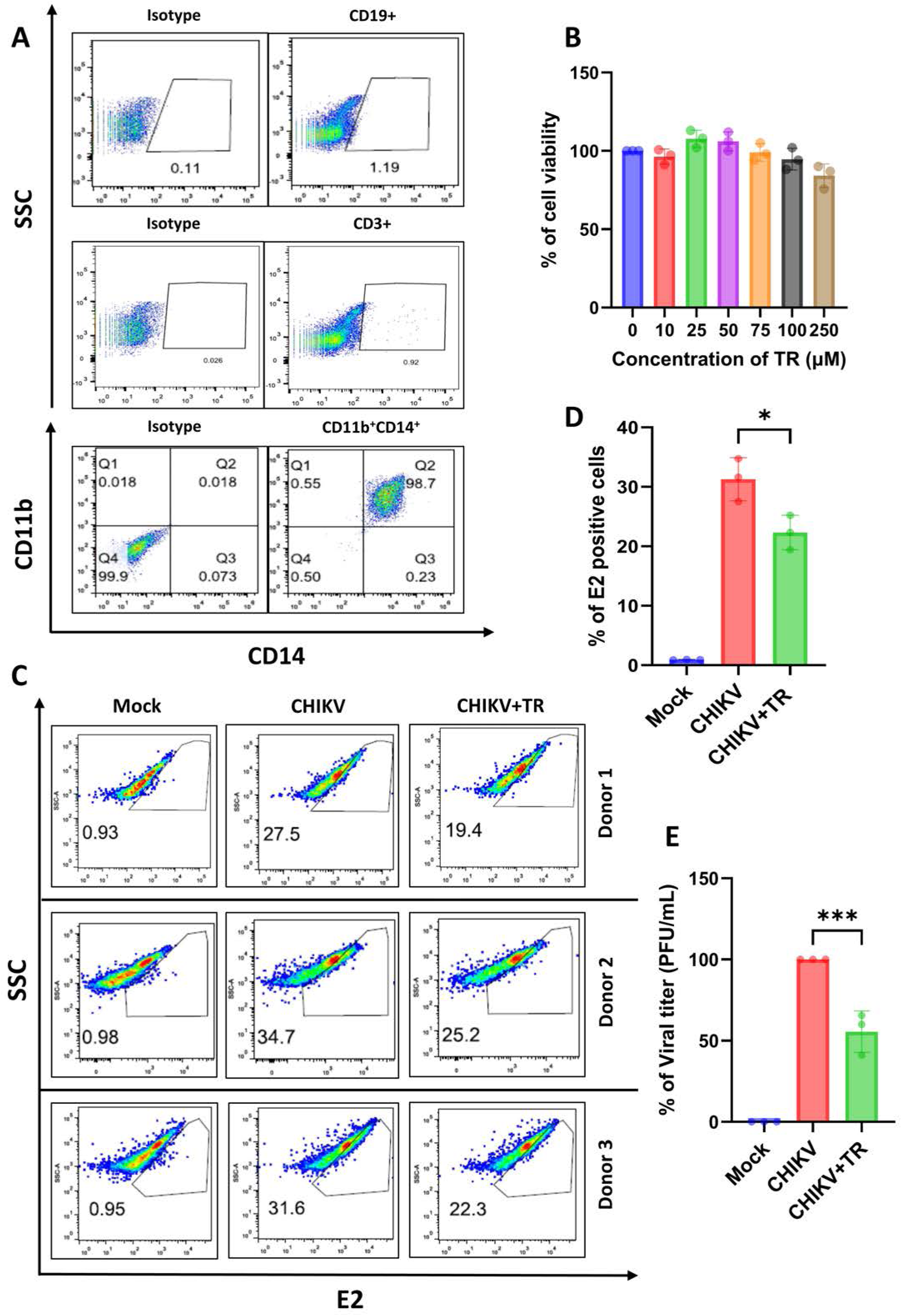
TR perturbs CHIKV infection in the hPBMC-derived monocyte-macrophage populations *ex vivo*. The human PBMCs were isolated from the blood samples of three healthy donors and subjected to immunophenotyping analysis of adherent and non-adherent populations. (A) Flow cytometry dot plots showing the percentages of B cells (CD19), T cells (CD3) and CD14^+^CD11b^+^ monocyte-macrophage cells from adherent hPBMCs. (B) Bar diagram showing the viability of CD14^+^CD11b^+^ monocyte-macrophage cells in presence of different concentrations of TR, obtained via MTT assay. (C) The hPBMC-derived monocyte-macrophage cells were infected with CHIKV-IS at 5 MOI and treated with TR. At 12 hpi, cells and supernatants were collected for downstream experiments. Flow cytometry-based dot plot analysis showing the percentage of viral E2-positive hPBMC-derived monocyte-macrophage populations in mock, CHIKV-infected and TR-treated samples. (D) Bar diagram representing the percent positive cells for CHIKV-E2 protein, obtained by flow cytometry. (E) The bar diagram depicts the % viral titer of the supernatants from CHIKV-infected and treated CD14^+^CD11b^+^ monocyte-macrophage cells. All the experiments were carried out in triplicate and data are shown as mean ± SD; p < 0.05 was considered statistically significant.

## DISCUSSION

Research on the antiviral compounds against alphaviruses, including CHIKV, remains mostly at the preliminary stage, lacking pre-clinical and clinical potencies (37), underscoring the unmet need for the development and evaluation of novel therapeutic approaches. Previous studies have demonstrated the broad-spectrum antiviral activity of retinoids against diverse viral families; however, the efficacy against alphaviruses has not been explored yet. This study reports the first evidence of inhibitory potential of ATRA or TR against CHIKV infection.

In the current investigation, TR was able to abrogate CHIKV infection significantly in Vero as well as physiologically relevant muscle cells, C2C12 with drastic reduction in viral RNA, proteins and release of infectious virus particles in a dose-dependent manner. The IC_50_ of TR against CHIKV was estimated to be 20.05 µM and 19.71 µM in the Vero and C2C12 cells, respectively along with good selectivity indices (8.34 in Vero and 9.61 in C2C12). Additionally, the the drug was effective during entry and post-entry phases of infection and interfered in the early stages of CHIKV life cycle, post-infection. TR also showed remarkable virucidal effect against CHIKV. Further, bulk RNA-seq identified a series of genes and cellular pathways which were altered upon CHIKV infection and drug treatment, compared to uninfected condition. Interestingly, the RAR receptors belonging to the nuclear receptor family were found to be key mediators for the anti-CHIKV action of TR. Inhibition of RAR-β by specific antagonist impaired the efficacy of the drug, suggesting the possible role of RAR-β in mediating the antiviral potential of TR against CHIKV. Furthermore, TR modulated CHIKV-induced inflammatory response *in vitro* by reducing the activated MAPKs, NF-κB and proinflammatory cytokines. Importantly, the antiviral efficacy of the drug was also validated *in vivo* using mice model. TR could protect CHIKV-infected C57BL/6 mice by diminishing viral burden leading to minimized disease score and better survival. Additionally, the abrogation of CHIKV infection by TR was also evident *ex vivo* in the hPBMC-derived monocyte-macrophags, supporting its preclinical efficacy.

Several studies reported earlier the broad-spectrum antiviral activity of various retinoids. For instance, pre-treatment with physiological concentrations of vitamin A or its metabolite ATRA suppressed HIV-1 replication in primary monocyte derived macrophages (MDMs) as well as HIV-1 p24 antigen production in THP-1 monocytes; however, the antiviral properties were ineffective upon retinoid-treatment post-infection (38). Both ATRA and 9cis-RA were found to inhibit Measles virus (MV) replication through the modulation of the RIG-I and nuclear retinoid receptor signaling pathways (19, 31). Similarly, another study has shown *in vitro* inhibition of Mumps virus (MuV) by Retinol and ATRA (39). Additionally, anti-hepatitis B virus (HBV) a tivity of natural and synthetic retinoids has been reported in primary human hepatocytes (PHHs) (22). Further, a drug repurposing screen of 1,018 FDA-approved compounds had also identified few retinoic acid receptor (RAR) agonists (Tretinoin, Acitretin, Adapalene and Tazarotene) as potent HBV inhibitors (23). Notably, the acyclic retinoid, Peretinoin demonstrated strongest antiviral potential against HCV infection, abrogating replication and infectious virus release by 80–90% without affecting the viral assembly (25). Another study revealed that ATRA binding to cellular retinoic acid binding protein 2 (CRABP2) potently abrogated HCV infection by blocking lipid droplets (LDs) accumulation, whereas engagement of CRABP1 by ATRA exerted a proviral effect by abundant accumulation of LDs (40). Furthermore, ATRA was also found to exhibit antiviral effects against various strains of SARS-CoV-2 by targeting the 3CLpro Protease Activity (27) as well as disrupting the spike-mediated cellular entry (28). This study highlights the anti-CHIKV property of TR, which is comparable with the previous observations.

Pre-treatment with the drug had no effect on viral particle formation, whereas reduction in viral progeny release in case of during-treatment indicates that TR may affect the viral attachment or entry process. Interestingly, highest inhibition was observed when the treatment was applied post-infection, after the entry of the virus inside the host cell. Thus, it can be speculated that TR likely targets multiple phases of CHIKV life cycle to modulate the infection process. Additionally, the virucidal assay revealed that TR might directly impair viral particles, thereby hampering viral attachment or the entry process, which requires further investigations. Although the mechanistic insights of retinoids against some viruses have slightly been explored, the in-depth molecular mechanisms of action as well as *in vivo* efficacy are majorly unknown.

Retinoids regulate the transcription of numerous retinoid responsive genes by activating the nuclear receptor complex, composed of the Retinoic Acid Receptor (RAR) and Retinoid X Receptor (RXR) heterodimer (41), which binds to the Retinoic Acid Response Elements (RARE) on the promoters of target genes. This work demonstrates that nuclear retinoid receptor signaling is central to the antiviral efficacy of TR against CHIKV. The transcriptomics analysis revealed that Signaling by nuclear receptors was among the key pathways upregulated upon addition of TR to the CHIKV infected cells. Although more than one nuclear receptor may be involved, a crucial role for RAR-β was observed. To support this, specific antagonists of RAR-α or RAR-β or RAR-γ (RO, LE and LY) were used and inhibition of RAR-β significantly antagonized the the anti-CHIKV action of TR, suggesting its possible role in mediating the antiviral effect. Apart from the nuclear receptor signaling, several other pathways were found to be modulated upon infection and drug-treatment in C2C12 cells, such as Transport of small molecules, Biological oxidations, Innate immune system, Kinesins, COP I-dependent Golgi to ER retrograde traffic and Golgi to ER retrograde transport. Further research may aid in actual mechanistic details of the antiviral features of TR, underlying the role of novel host factors associated with these cellular pathways.

Regulation of the pattern recognition receptor RIG-I and IFN regulatory factor 1 (IRF-1) by retinoids is a crucial factor to enhance innate immune responses and antimicrobial defense (23, 31). Several studies have deciphered the antiviral effect of retinoic acids against HIV, HCV, MeV and MuV through the activation of innate immune pathways and RIG-I signaling (20, 31, 42, 43). However, in the current study, the anti-CHIKV potential of TR was found to be RIG-I independent, as demonstrated by using the Huh-7/7.5 cell culture model system. In this context, the roles of the genes associated with the interferon signaling and antiviral response can be investigated further. Inflammation induced by CHIKV infection is the major cause of its morbidity. MAPKs are activated in response to CHIKV infection (33) and inhibition of the phosphorylated MAPKs resulted in significant abrogation of viral infection (34). TR was able to reduce the key proinflammatory mediators and transcription factors (p38, ERK1/2, SAPK/JNK, NF-κB, IRF3) in CHIKV-infected murine macrophages, establishing its anti-inflammatory role. This was also supported by the reduction in the levels of cytokines like TNF-α, MCP-1 and IL-6, which are central to the inflammatory process. In agreement to this, the drug was able to manage the CHIKV-induced immune response following infection.

The *in vitro* findings were corroborated further in the C57BL/6 mouse model. Viral burden and proteins were reduced significantly in the tissues of the infected mice along with reduced clinical scores upon oral administration of TR (5 mg/kg). The observed *in vivo* efficacy of TR against CHIKV infection at an acceptable human equivalent dose suggests its suitability for repurposing against this disease. Although extrapolation from pre-clinical models to clinical settings may not be perfectly linear, the anti-CHIKV property of the drug was also evaluated in hPBMC-derived monocyte-macrophage populations. Reduction of viral titer and viral protein levels in the CHIKV-infected human PBMCs upon TR-treatment support its suitability for further clinical application.

Despite the broad antiviral potential of retinoic acids, the pharmacological limitations including poor solubility (44), weak photostability (45), and toxicity (46, 47) affect their therapeutic efficacy. Owing to the challenges, advanced retinoid delivery formulations based on gels, liposomes, microparticles, nanoparticle are under investigation for improving solubilization, reducing toxicity and sustaining release towards enhanced translational potential (48).

In summary, this study demonstrated the anti-CHIKV efficacy of TR *in vitro* in multiple cell lines, *in vivo* in C57BL/6 mice and, *ex vivo* in hPBMC-derived monocyte-macrophage populations, highlighting its repurposing potential towards future therapeutics against this viral infection.

## MATERIALS AND METHODS

### Cells and Virus

The Vero (African green monkey kidney epithelial cells), Huh7 (Human hepatocellular carcinoma cells) and RAW 264.7 (mouse monocyte/macrophage) cells were obtained from the National Centre for Cell Science (NCCS), Pune, India. The mouse myoblast cell line C2C12 was generously gifted by Dr. Amresh C Panda, Institute of Life Sciences (ILS), Bhubaneswar, India. Huh7.5 cells were provided by Apath, LLC (New York, NY, USA). Vero and C2C12 cells were cultured in Dulbecco’s Modified Eagle Medium (DMEM; Himedia, India) supplemented with 10% (for Vero) or 15% (for C2C12) fetal bovine serum (FBS; Gibco, USA) along with gentamicin (Gibco, USA) and penicillin-streptomycin (Himedia, India). For the Huh7 and Huh7.5 cells, non-essential amino acids (Gibco, USA) were additionally complemented to the DMEM media (previously described) for optimal cell growth. The RAW 264.7 cells were maintained in Roswell Park Memorial Institute (RPMI) 1640 Medium (Gibco, USA) supplemented with 10% FBS, gentamicin and penicillin-streptomycin. All the cells were grown at 37°C with 5% CO_2_ inside a humidified incubator.

The CHIKV prototype strain (PS, accession no. AF369024.2) was a kind gift from Dr. M.M. Parida, Defence Research and Development Establishment (DRDE), Gwalior, India. The Indian outbreak strain of CHIKV (IS; accession no. PP349434) was also used in this study. All the experimental protocols involving CHIKV were conducted according to the guidelines of Institutional Biosafety Committee (IBSC).

### Antibodies and inhibitors

The CHIKV-E2 monoclonal antibody (mAb) was kindly provided by Dr. M.M. Parida (DRDE, Gwalior, India). The CHIKV-nsP2 mAb was developed and characterized by our group (49). The p-P38, P38, p-ERK1/2, ERK1/2, p-SAPK/JNK, SAPK/JNK, p-NF-κB, NF-κB, p-IRF3, IRF3, β-Actin, RIG-I and F4/80 antibodies were purchased from Cell Signaling Technology (CST; USA). The Glyceraldehyde 3-phosphate dehydrogenase (GAPDH) antibody was procured from ABGENEX Pvt. Ltd. (India). HRP-conjugated anti-mouse and rabbit secondary antibodies were purchased from Promega, USA. Tretinoin (TR), Ro 41-5253 (RO), LE135 (LE) and LY2955303 (LY) were obtained from Sigma Aldrich (USA).

### Cytotoxicity assay

To determine the cytotoxicity of TR, RO, LE and LY, 3-(4,5-dimethyl-2-thiazolyl)-2,5 diphenyl-2H-tetrazolium bromide (MTT) assay was conducted using the EZcount MTT cell assay kit (Himedia, India) as per the manufacturer’s protocol. Briefly, cells were seeded in 96-well plates, one day before the experiment. At 80% confluency, the cells were treated with increasing concentrations of drug/inhibitors along with dimethyl sulfoxide (DMSO) as the reagent control. After 16-24 hours (h) post-treatment (time of incubation varied based on cell lines used), MTT solution (5 mg/mL) was added to the cells, followed by incubation of the culture plates at 37°C for 2 h. Next, the medium was removed and solubilization solution (100 μL/well) was added to dissolve the formazan crystals. Finally, the absorbance was measured at 570 nm using a multimode plate reader (PerkinElmer, MA) and the percentage of metabolically active cells was compared with the control cells to estimate cell viability as described earlier (50).

### Virus infection

The Vero, Huh7 and Huh7.5 cells were infected with 0.1 multiplicity of infection (MOI) and the C2C12 cells were infected with 0.05 MOI of CHIKV-PS, according to the protocol mentioned before (50). In case of RAW 264.7 cells, infection was carried out using CHIKV-IS at an MOI of 5, as reported earlier (51). After 90 minutes (min) incubation period at 37°C with shaking at every 10 to 15 min intervals, the inoculum was replaced with different concentrations of TR, diluted in fresh medium. The mock, infected and treated cells were examined under a microscope (20X magnification) to observe the cytopathic effect (CPE). Next, the culture supernatants and cells were harvested at different hours post-infection (hpi) for downstream experiments.

### Plaque assay

Plaque assay was performed to quantify the viral titer, according to the protocol described previously (52). In brief, the Vero cells were infected with different dilutions of the cell supernatants or tissue homogenates of mock, infected and drug-treated samples for 90 min. Then, the cells were washed with 1X phosphate-buffered saline (PBS) and subsequently overlaid with 2% methylcellulose (Sigma, USA), followed by incubation at 37°C for 4-5 days. Next, the cells were fixed with 8% formaldehyde and stained with crystal violet to count the plaques and viral titers were exhibited as plaque-forming units per milliliter (PFU/mL).

### Confocal microscopy

Immunofluorescence imaging was performed as mentioned before (53). The Vero and C2C12 cells were seeded on coverslips in 6-well culture plates. Upon reaching 60% confluency, the cells were infected with CHIKV, followed by drug treatment. The cells were then fixed with 4% paraformaldehyde (PFA) at 16-18 hpi (16 hpi in case of C2C12 and 18 hpi for the Vero cells) and subsequently incubated with CHIKV-E2 primary antibody (1:750 dilution). After 1 h, the cells were washed with 1X PBS and secondary anti-mouse Alexa Fluor (AF) 488 antibody (Invitrogen, USA) was added at 1:1000 dilution for 45 min. Next. the cells were stained with 4′,6-diamidino-2-phenylindole (DAPI; Life Technologies, Invitrogen, USA) and coverslips were mounted with the antifade reagent (Invitrogen, USA) to prevent photobleaching. Finally, fluorescence microscopic images were captured using the Leica TCS SP5 confocal microscope (Leica Microsystems, Germany) and the images were analysed using the ImageJ software.

### qRT-PCR

Total cellular RNA was extracted using the Trizol reagent (Ambion, USA) from the mock, infected and treated cells as reported earlier (54). An equal amount of RNA was reverse-transcribed into cDNA using using the PrimeScript™ 1st strand cDNA Synthesis Kit (Takara Bio, USA). For qRT-PCR, CHIKV E1 and nsP2 genes-specific primers were used along with GAPDH or β-Actin as endogenous controls (50). Similarly, equal volume of culture supernatants or mice serum samples were used for viral RNA extraction using the QIAamp Viral RNA mini kit (Qiagen, Germany), following the manufacturer’s instructions. An equal volume of viral RNA was used to synthesize cDNA and E1 gene was amplified by qRT-PCR with equivalent volume of cDNA. CHIKV RNA copy numbers were calculated from the corresponding C_t_ values, using a standard curve (52).

### Western blot

Western blot analysis was executed according to the procedure described before (55). Briefly, cells were harvested at different time points and lysed subsequently using equal volume of Radio Immuno Precipitation Assay (RIPA) lysis buffer containing Protease and Phosphatase inhibitor cocktails (Roche, Germany). Snap-frozen mock, infected and infected/drug-treated mouse tissues (brain and muscle) were homogenized and lysed in RIPA buffer. The protein lysates were quantified using the Pierce™ BCA Protein Assay Kit (Thermo Fisher Scientific, USA). Next, equal concentrations of protein were separated on 10% SDS-polyacrylamide gel and transferred onto a polyvinylidene difluoride membrane (PVDF; Millipore, USA), followed by blocking with 3% BSA (Sigma-Aldrich, USA). The membrane was then probed using primary antibodies with dilutions as per the manufacturer’s recommendation. GAPDH and Actin antibodies were used as loading controls. After primary antibody incubation, corresponding anti-Mouse or Rabbit HRP conjugated secondary antibodies were added to the blots. Finally, the blots were developed using the Immobilon Western Chemiluminescent HRP substrate (Millipore, USA) in the ChemiDoc MP Imaging System (Bio-Rad Laboratories, USA), and band intensities were quantified using the ImageJ software.

### Flow cytometry

Flow cytometry-based analysis was carried out as reported earlier (34). Briefly, mock, infected and drug-treated cells were harvested and fixed in 4% PFA for 10 min at room temperature (RT), followed by washing with 1X PBS and re-suspension in FACS buffer (1X PBS, 1% BSA, 0.01% NaN_3_). For intracellular staining (ICS), the fixed cells were permeabilized with permeabilization buffer (1X PBS, 0.5% BSA, 0.1% Saponin and 0.01% NaN_3_), followed by blocking with blocking buffer (1X PBS, 1% BSA, 0.1% Saponin and 0.01% NaN_3_) for 30 min at RT. Next, the cells were sequentially incubated with CHIKV-E2 primary and anti-mouse AF-647 conjugated secondary antibodies, diluted in permeabilization buffer for 30 min at RT. Finally, the cells were washed with permeabilization buffer and suspended in FACS buffer and approximately 1 × 10^4^ cells were acquired per sample using the BD LSRFortessa™ Flow cytometer (BD Biosciences, USA) and analyzed by the FlowJo™ software (BD Biosciences, USA).

### Treatment at different phases of viral infection (Pre, during and post infection)

In order to decipher the impact of the drug on CHIKV infection cycle, TR (50 μM) was administered at different phases of infection (pre, during and post treatment) in both Vero and C2C12 cells, as described earlier (56).

In pre-treatment, cells were first pre-treated with TR for 3 h, followed by washing with 1X PBS and then CHIKV infection was carried out for 90 min. Next, the cells were washed again and fresh media was added without any drug.

For the during-treatment condition, cells were simultaneously incubated with the virus inoculum and TR for 90 min at 37°C. The mixture was then removed and the cells were washed with 1X PBS, followed by the addition of fresh media without TR.

In case of the post-treatment, cells were first infected with CHIKV for 90 min. The infected cells were then washed with 1X PBS and drug-containing media was added to the cells.

Finally, supernatants from the Vero and C2C12 cells were collected at 18 hpi and 16 hpi, respectively and subjected to plaque assay.

### Virucidal Assay

To assess the extracellular effect of TR on CHIKV, virucidal assay was performed as mentioned previously (57). CHIKV (0.1 MOI) was incubated with 50 µM of TR for 30 min in serum free media at 37°C. Then, plaque assay was performed with the untreated and drug-treated viral inoculums to determine the residual infectivity.

### Time-of-addition experiment

To estimate the effectiveness of TR on CHIKV replication cycle post-infection, time-of-addition experiment was carried out, as mentioned before (53). Briefly, TR (50 μM) was added to both the Vero and C2C12 cells at 0, 2, 4, 6, 8, 10, 12, 14 and 16 hpi, following infection with CHIKV. The culture supernatants from all the time points were collected at 16-18 hpi (16 hpi for C2C12 and 18 hpi for Vero cells) and viral titer was determined by plaque assay.

### Library preparation for RNA Sequencing (RNA-seq)

The RNA-seq library preparation was performed as described earlier with little modifications (58, 59). In brief, total RNA of each C2C12 cell sample (Mock, Mock + TR, CHIKV Infected and CHIKV Infected + TR) was extracted using the Trizol reagent (Ambion, USA) and RNA quantity and purity were assessed through NanoDrop 2000 Spectrophotometer (Thermo Scientific, USA) and 4200 TapeStation system (Agilent Technologies, USA), respectively. Next, 1μg of total RNA with RNA integrity number (RIN) ≥ 9 was taken to isolate mRNA using magnetic beads and an mRNA isolation kit (NEBNext Poly(A) mRNA Magnetic Isolation Module, New England Biolabs, USA). For library preparation, NEBNext Ultra II RNA Library Prep Kit for Illumina was used as per the manufacturer’s instructions. The prepared libraries of three biological replicates of each sample were then quantified by the Qubit 4 Fluorometer (Thermo Fisher Scientific, USA) and Tapestation Bioanalyzer was used to examine the fragment sizes. The cDNA samples were denatured using 0.2 N NaOH for 5 min at RT. Further, to neutralize the impact of NaOH, 10 mM Tris-Cl (pH-8.5) was added. Finally, the 4 nM cDNA libraries were diluted with HT1 buffer for a final loading concentration of 400 pM, pooled and pair-end sequenced on the NovaSeq 6000 Sequencing System (Illumina, USA) at the in-house Sequencing facility of ILS.

### RNA-seq data analysis

The short reads obtained from the sequencing were screened to remove low-quality reads and adapter sequences using Trimmomatic v 0.39 (60). The clean reads were mapped onto the CDS sequences annotated for *Mus musculus*, available in the Ensembl database (https://ftp.ensembl.org/pub/release-114/fasta/mus_musculus/). Normalization of gene counts and differential expression estimations were completed in R studio (http://www.rstudio.com/) version 4.4.2 using DESeq2 package. Significantly differentially expressed genes (DEGs) were analysed (adjusted p-value < 0.05 and |log_2_FC ≥1|) across different experimental conditions. The Reactome pathway enrichment using the enrichPathway function and Gene Set Enrichment Analysis (GSEA) were carried out using the significant DEGs (61). Pathways with a p-value and q-value below 0.05 were considered significantly enriched, providing insights into the biological processes altered by CHIKV infection and TR treatment. A total of 240,750,092 reads were obtained after RNA-seq and were screened for quality and adapter trimming to retain 231,195,532 reads (details have been provided in supplementary Table S1).

### siRNA transfection

siRNA based gene knockdown was performed as reported earlier (57). Monolayers of Huh7 cells with 60% confluency were transfected with 50 pM of RIG-I siRNA (sc-61480, Santa Cruz Biotechnology, USA) using Lipofectamine-3000 reagent (Invitrogen, USA) in Opti-MEM Reduced Serum Medium (Gibco, USA) as per the manufacturer’s instruction. At 36 h post-transfection (hpt), the cells were infected with CHIKV at an MOI of 0.1 as described above and TR was added post infection. The supernatants were harvested at 24 hpi for plaque assay.

### Sandwich enzyme-linked immunosorbent assay (ELISA)

The RAW 264.7 cells were infected with CHIKV and treated with 50 and 100 μM of TR. The cell-free culture supernatants were harvested at 8 hpi and the levels of different cytokines, viz. tumor necrosis factor-alpha (TNF-α), interleukin-6 (IL-6) and monocyte chemoattractant protein 1 (MCP-1) were quantified by sandwich ELISA using the BD OptEIA™ Sandwich ELISA kit (BD biosciences, USA) according to the previously described protocol (62). The concentrations of cytokines in the test samples were calculated with respect to corresponding standard curves constructed using different known concentrations of recombinant cytokines in pg/mL (51). The cytokine concentrations were measured at 450 nm using the Epoch 2 (BioTek, USA) microplate reader.

### Animal studies

Animal experiments were executed following the guidelines of the Committee for the Purpose of Control and Supervision of Experiments on Animals (CPCSEA) of India along with the approval of the Institutional Animal Ethics Committee (ILS/IAEC-275-AH/MAY-22). 10-12 days old C57BL/6 mice were infected subcutaneously with 10^6^ PFU of CHIKV-PS at the flank region of the right hind limb while the control mice (mock) were injected with serum-free media as described earlier (50). After 3 hpi, TR (5mg/kg) was administered orally to the treated group of mice (n=3) and drug-treatment was continued at an interval of 24 h up to 4 days post infection (dpi). Solvent was provided to the mock and infection-control groups of mice (*n* = 3). On 5 dpi, the mice were sacrificed, sera were isolated from the blood and different tissues (quadriceps muscle and brain) were collected and snap-frozen in liquid nitrogen for Western blotting or stored in 10% formalin for histological and immunohistochemical studies. Viral load in the serum was assessed by qRT-PCR and the serum TNF level was quantified by ELISA-based cytokine assay. Further, an equal amount of tissue from each group (mock, infected and treated) was homogenized in serum-free media (DMEM), followed by syringe filtration using a 0.22-μm membrane. The solutions were then centrifuged and the supernatants were collected for plaque assay to determine the viral burden in the tissues as mentioned before (52). For the clinical score and survival curve analyses, similar protocol for infection and treatment was followed as mentioned above (n=6 mice for all three groups) and the drug was administered from 0-10 dpi. The mice were thoroughly monitored on regular basis for the development of disease symptoms and mortality. The clinical score of each mouse was tabulated regularly according to onset of disease symptoms (no symptoms: 0; hunchback: 1; single hind limb paralysis: 2; both hind limb paralysis: 3; loss of mobility: 4 and Death: 5). Finally, the survival curve was made on the basis of mortality (50, 52).

### Histology and Immunohistochemistry

The histopathological investigations were carried out according to the protocol mentioned before (50, 63). In brief, formalin-fixed tissue samples were serially dehydrated and embedded in paraffin wax and paraffin blocks were cut into sections of 5 μM thickness, using a rotary microtome (Leica Biosystems). Next, the sections were stained with haematoxylin and eosin (H&E; Himedia, India) and histological changes were visualized using a light microscope (Zeiss Vert.A1, Germany). For Immunohistochemistry, the tissue sections on glass slides were incubated with the primary CHIKV-E2 or F4/80 antibodies for overnight at 4°C, followed by washing with 1X PBS. Next, secondary anti-mouse AF 488 (or 594) and anti-rabbit AF 647 antibodies (Invitrogen, USA) were added to the sections for 45 min at RT inside a dark and humidified chamber. The slides were then stained with DAPI (Life Technologies, Invitrogen, USA) and coverslips were mounted using the mounting reagent. Finally, the slides were observed under the Apotome fluorescence microscope (Zeiss, Germany).

### Isolation of human peripheral blood mononuclear cells (hPBMCs)

Isolation of hPBMCs was performed from the blood samples of three healthy donors as described previously (55). Human blood was collected, following the guidelines of the Institutional Ethics Committee, NISER, Bhubaneswar (NISER/IEC/2022-04). Briefly, 30 mL blood was collected from each of the healthy individuals and subjected to Hi-Sep LSM (HiMedia, India) based density gradient-centrifugation to isolate PBMCs (52). The extracted PBMCs were cultured in RPMI 1640 Medium (Gibco, USA) supplemented with 10% FBS and antibiotic-antimycotic solution for 5 days. Next, hPBMC-derived adherent cells were undergone immunophenotyping analysis and found to be enriched mostly with CD14^+^CD11b^+^ monocyte-macrophage lineages.

### CHIKV infection in hPBMC-derived monocyte-macrophage cells and drug treatment

At first, the cytotoxicity of TR for the adherent hPBMCs (CD14^+^/CD11b^+^ cells) was determined via MTT assay as mentioned above. The adherent cells were then infected with CHIKV-IS at 5 MOI for 2 h and treated subsequently with TR (100 μM). The infected and treated cells as well as supernatants were collected at 12 hpi and subjected to flow cytometry based intracellular staining for the detection of viral protein E2 or plaque assay to estimate viral titer (52, 53).

### Statistical analysis

All the statistical analyses were performed using the GraphPad Prism version 8 software. Group comparisons were conducted using the one-way or two-way analysis of variance (ANOVA), depending on the experimental design. For the comparison of two groups, unpaired two-tailed Student’s *t*-test was performed. Data of three independent experiments are shown as mean ± standard deviation (SD) unless stated otherwise. A p-value of < 0.05 was considered statistically significant (denoted by asterisks, with *p <0.05, **p ≤0.01, ***p ≤0.001 and ****p ≤0.0001; ns: non-significant).

## Data availability

The data that support the findings of this study have been included in the manuscript. The RNA-Seq data discussed in this manuscript have been deposited in the Sequencing Read Archive (SRA) at the National Center for Biotechnology Information (https://www.ncbi.nlm.nih.gov/) under the BioProject accession number: PRJNA1309963 (https://dataview.ncbi.nlm.nih.gov/object/PRJNA1309963?reviewer=s4t81itg020t8cm72r1abk39rf).

## ACKNOWLEDGMENTS

This work was funded partly by the Department of Biotechnology (DBT), Ministry of Science and Technology, Government of India (Grant No. BT/PR42322/TRM/120/525/2021) and partly by the ILS Core Fund provided by the DBT. Soumyajit Ghosh was supported by a Senior Research Fellowship (Award Letter No. VIR/Fellowship/2/2022-ECD-I) from the Indian Council of Medical Research (ICMR), Government of India. The funding agencies had no role in the study design, experiments, analyses, data interpretation, manuscript writing or in the decision to publish the results.

We are grateful to Dr. M. M. Parida, DRDE, Gwalior, India, for kindly providing the CHIKV-PS strain and CHIKV-E2 monoclonal antibody. We sincerely acknowledge the Central Instrumentation Facility (CIF), Core Sequencing facility and the Animal House facility of ILS, Bhubaneswar. We would also like to thank Surajit Gandhi, Kaushik Sen, Supriya Suman Keshry, Koustav Chatterjee, Ankita Datey, Kshyama Subhadarsini Tung and Anjali Girish for their assistance during the experiments and valuable suggestions. The schematic representations of the RNA-Seq workflow and animal experiments have been created using BioRender.com.

## AUTHOR CONTRIBUTIONS

SG and SoC conceived the idea, designed the experiments, and analyzed and interpreted the results. SoC, SuC and SP contributed reagents and other necessary resources. SG, CM, AM, TM, BB, SD, UG and RRJ carried out the wet lab experiments. PK and SP performed the *in silico* experimentation and data analysis. SG prepared the figures, and wrote and edited the manuscript. SoC reviewed the manuscript and approved the final version.

